# OpenStats: A Robust and Scalable Software Package for Reproducible Analysis of High-Throughput Phenotypic Data

**DOI:** 10.1101/2020.05.13.091157

**Authors:** Hamed Haselimashhadi, Jeremy C Mason, Ann-Marie Mallon, Damian Smedley, Terrence F Meehan, Helen Parkinson

## Abstract

Reproducibility in the statistical analyses of data from high-throughput phenotyping screens requires a robust and reliable analysis foundation that allows modelling of different possible statistical scenarios. Regular challenges are scalability and extensibility of the analysis software. In this manuscript, we describe OpenStats, a freely available software package that addresses these challenges. We show the performance of the software in a high-throughput phenomic pipeline in the International Mouse Phenotyping Consortium (IMPC) and compare the agreement of the results with the most similar implementation in the literature. OpenStats has significant improvements in speed and scalability compared to existing software packages including a 13-fold improvement in computational time to the current production analysis pipeline in the IMPC. Reduced complexity also promotes FAIR data analysis by providing transparency and benefiting other groups in reproducing and re-usability of the statistical methods and results. OpenStats is freely available under a Creative Commons license at www.bioconductor.org/packages/OpenStats.

## Introduction

Statistics is the main inferential tool used in science and medicine to extract information from data. It provides a set of proven steps for drawing conclusions and making decisions in spite of the uncertainty inherent in any data, which are unavoidable due to biological variation as well as the constraints of cost, time, and measurement precision. The inference made from the data is subject to reproducibility in the analysis requiring precise, transparent, comprehensive and well-documented rules to prevent unreliable, costly and even invalid results ^1^. Reviewing the literature shows that reproducibility in the data and analysis is the subject of an extensive range of publications in different areas of science, e.g., life science, bioscience, medical and pharmaceutical science and translational science ^2–6^. Studies have shown irreproducibility of results is often due to poor documentation of the statistical method ^7–9^. This is especially critical for the high-throughput phenomic screening when tens of thousands of data points are generated and analysed.

The International Mouse Phenotyping Consortium (IMPC www.mousephenotype.com) is a G-7 recognised global research infrastructure dedicated to generating and characterising a knockout mouse line for every protein-coding gene ^10–13^. Currently, in its 8^th^ year, the IMPC has phenotyped over 188K + knockout and 69K + control mice across 13 research centres from 9 countries in 3 continents (data release 11, February 2020, www.mousephenotype.org/data/release). These centres adhere to a set of standardised phenotype assays defined in the International Mouse Phenotyping Resource of Standardised Screens (IMPReSS www.mousephenotype.org/impress) that represents over 496 (116 IMPC specific) procedures and 10,266 (3,054 IMPC specific) parameters measured on mice. As part of the operating design protocol, critical factors that can impact data collection such as reagent type or equipment are captured as mandatory metadata. Phenotype data is then centrally collected and quality controlled by trained professionals before being released for statistical analyses.

The data is then processed by PhenStat ^14^, a freely-available R ^15^ package that provides a set of statistical methods for the identification of genotype to phenotype associations by comparing mutants to controls ^16^. PhenStat imposes the same statistical model on the entire continuous data regardless of the nature of the original measurements. That is, the continuous measurements are analysed using a Linear Mixed Model (LMM) ^17^ under the following setting for the fixed effects,

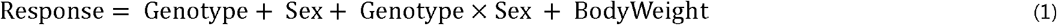

and Batch (defined as the date of measurement) in the random effect. For the cases where LMM fails, PhenStat proceeds to an alternative method called Reference Range Plus (RR+) ^16^. The RR+ method relies on an initial setting of a quantile, default 95% in PhenStat, to form the initial classes (Low/Normal/High) that discretises a continuous response based on the control, wild type (WT), mice population. Mutants are then stratified into the (Low-Normal versus High) and similarly to (Low versus Normal-High) classes. One can present the RR+ model as below,

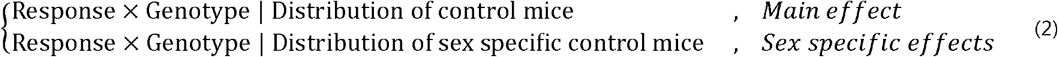

where (‘|’) and (‘x’) J represent the *conditional operation* and *interaction* respectively. Fisher’s Exact Test is applied to each contingency table to test the hypothesis of the independence of rows (discretised response) and columns (genotype). The categorical data in the IMPC are analysed using Fisher’s Exact test for combined sexes as well as the individual sexes under the model below,

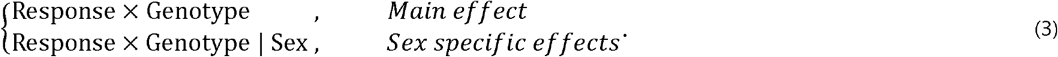

Besides the advantage of providing a reliable analysis pipeline by PhenStat, a number of limitations arise with the scalability of the input data flow and the diversity of scenarios that can be handled by the statistical software in high-throughput screening pipelines. For instance, the internal optimisation of PhenStat for continuous measurement relies on the repetitive use of Likelihood Ratio Tests (LRT) for model selection that requires a predefined threshold (default 0.05 in PhenStat). This leads to a reduction in transparency of the statistical analysis in addition to an increase in the computational complexity of the analysis for large scale screening projects such as IMPC. There are also many instances of the data in IMPC with repetition in the measurement values that lead to a misleading inference from the current implementation of the RR+ in PhenStat. These issues, coupled with the ever-growing screens in the IMPC, especially ageing (e.g., longitudinal data), require scalable and more versatile statistical pipeline with user-defined models for different possible scenarios.

In this paper, we address the issues of scalability, extensibility, versatility, and efficiency in the current IMPC statistical pipeline implemented using the R package PhenStat by introducing a new package that we call ***OpenStats*** in the same development environment, R. The new software offers versatile modelling of high-throughput phenotypic data, such as modelling time dependency in data, for example, the longitudinal data in the IMPC ageing pipeline, with a focus on simplicity and efficiency. We assess the performance of OpenStats on the IMPC data including more than 2.5M datasets and analyses and compare the results with the current implementation of the IMPC stats pipeline. OpenStats is available from the Bioconductor repository (www.bioconductor.org/packages/OpenStats) that can be installed using the standard R workflow.

## Methods

The IMPC data are collected from 13 research centres around the world and consists of 61M + data points (Data release 11, February 2020) from different measurements on the control and mutant mice. The Genotype effect is measured on a set of knockout mice ^18^, typically 14 (7 males and 7 females), and a large group of controls, normally several hundred mice, spread over a moderately long period of time from months to years. Figure 1 shows the increase in the number of phenotyped mice lines (left) as well as the data points (right) that are collected along with the IMPC data releases starting from the first release in 2012 to the current release in 2020. This figure shows that on average the total number of the data points and the phenotyped lines between major data releases are increased by a factor of 21% and 23% respectively. Along with the scalability of the data, one challenge is the long term variability such as seasonal effects, changes in personnel and unknown time-dependent environmental factors in the IMPC data that is addressed by SoftWindowing method ^19^. However, an accurate solution to the estimation of the long term variation in SoftWindowing requires a precisely formed *initial model* that is fitted to the data. This motivates a reimplementation of the current methods with more versatility in the configuration and robustness in the implementation for the initial model.

**Figure 1.**
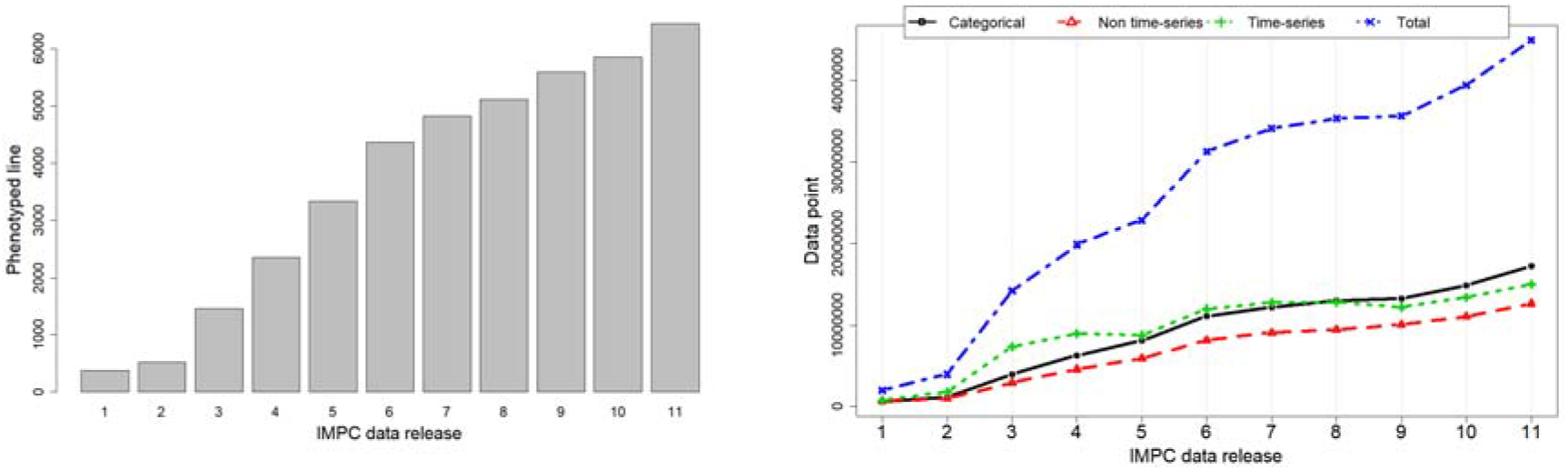
The increase in the total number of the IMPC mouse lines/data points along with the IMPC data releases from the first release in 2012 to the current release in 2020. (Left) The total number of IMPC phenotyped lines corresponding to the data releases. (Right) The overall increasing trend in the data points divided by the type of the data, non-time series (red), the time series (green), categorical (black) and total (blue) corresponding to the IMPC data releases. These plots show that on average the total number of data points and phenotyped mouse lines increase by a factor of 20% between IMPC data releases.

### Building block of the software

Figure 2 shows the four-layer structure of the software package namely, input data and model specification, data preparation, statistical analysis and reports.

**Figure 2.**
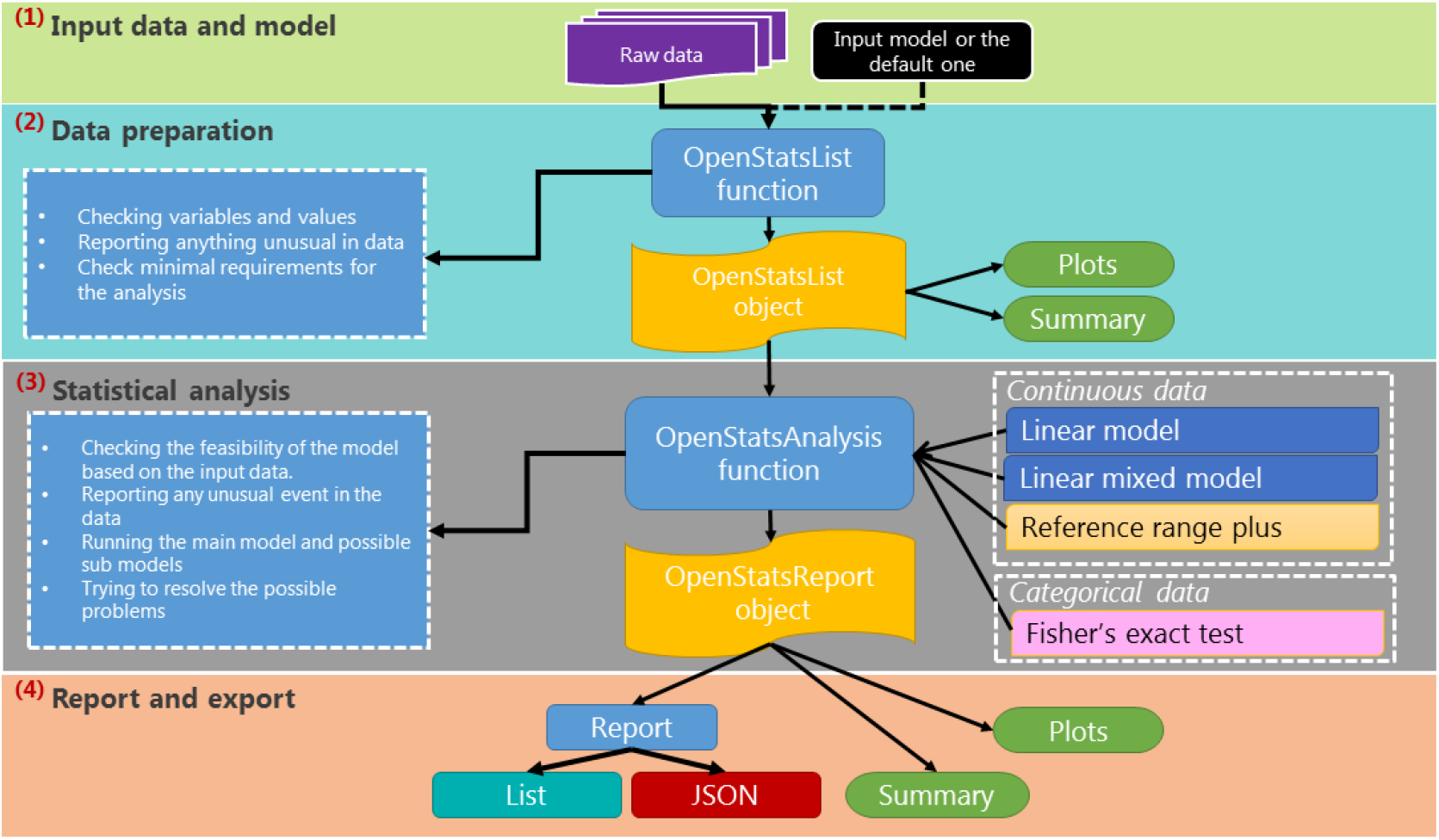
The schematic illustration of the OpenStats workflow. The OpenStats software is designed with a four-layer structure namely Input data and model specification, dataset processing and preparation, statistical analysis, and reporting/exporting the results.

The first layer, input data and model specification, data and the statistical model are mandatory, however, if the model is not specified, the default model is set to the same standard model ^20^ as PhenStat in Eq. (1), (2) and (3) corresponding to the type of input data.

The second layer, data preparation, the data and model terms are checked for missing data, redundancy, essential terms (such as genotype effect), mismatching between model terms and the input data, and normalising the different terminologies in the data e.g. sex, Sex, gender to a unified semantic (e.g. “Sex”). Furthermore, it allows basic standard operations such as missing specification, visualisations and summary via standard plot and summary functions.

The third layer, statistical analysis, is managed by the `OpenStatsAnalysis` function. This works as a hub for different statistical methods that can be selected using the “`method`” argument. Regardless of the chosen method, the function checks for the concordance of the input model and the underlying data and whether the type of data fits the chosen statistical model. This further checks for other input arguments to the function and reports any potential errors. For the implemented statistical frameworks, the statistical significance is assessed and several measures such as standardised effect sizes, confidence intervals, sex-specific effects (provided sex is included in the input data), summary statistics of the input data and several other measures are reported. Moreover, the `plot` and `summary` are available for all methods.

The final layer in the workflow, report and export, is managed by the function `OpenStatsReport` and allows the key elements of the analysis to be extracted in the form of either *list or JavaScript Object Notation* (JSON www.json.org). The output of the `OpenStatsReport` function has a schema that makes it versatile to be populated into databases or used by other software in a pipeline.

### Available statistical frameworks

Selection of the statistical method in the high-throughput screening pipelines that best fits the input data and the goal of the project is crucial otherwise can lead to misleading or weak evidence or increase the chance of producing Type I or Type II error in hypothesis testing ^21^. However, efficiency and simplicity are essential when hundreds of thousands of analyses are performed in the high-throughput screening pipelines such as in the IMPC. Below we describe the three main analysis frameworks that are implemented in the OpenStats software package.

### Nominal data

The majority of the nominal data in the IMPC measure the occurrence of a rare event such as *abnormal behaviour* or *absent/present of the tail* in the mice. To comply with the goal of the analysis, OpenStats applies Fisher’s Exact Test ^22–25^ with p-values computed by Monte Carlo simulation for larger than *2 by 2* contingency tables. Depending on the specification of the model, sub-tables such as male, female, lifestage (defined as Early/Late adult mice), male/female × LifeStage interactions etc. are also formed and tested. The confidence interval for the odds ratios of 2 × 2 tables and effect sizes, defined as the maximum percentage change from the corresponding contingency table, are estimated for all tables.

### Continuous data

#### Linear mixed model

The majority of the phenotypic data from the IMPC are continuous and analysed by performing the linear mixed model ^17,26^ with Genotype, Sex and Bodyweight in the covariates and Batch (as the date of measurement) in the random effect ^27,28^. OpenStats allows an open structure for the covariates, random effect and the further structures on the within/between-group variation. This allows modelling of complex data in the IMPC such as repeated measures by including custom covariates/random effects. To cope with the low *N*, typically 4-7 mutant animals per sex in high-throughput pipelines, OpenStats applies an optional forward/backward/stepwise ^29^ optimisation to all model terms on the basis of comparing mutual AICc, a version of Akaike information criterion (AIC) that has a correction for small sample sizes ^30^. Further to the initial model that is specified by the user, the sub-models are also fitted to the data for special purposes such as the detection of the sex specific (sexual dimorphism) effects. The confidence intervals, standard effect sizes ^31^ and standardised coefficients are also estimated for each possible sub-model. OpenStats further allows diagnosing the fitted model by providing visualisation tools and also reporting Shapiro-Wilk or Kolmogorov-Smirnov normality test statistics ^32,33^ for assessing the normality of the model residuals.

#### Reference Range Plus method

The Reference Range Plus (RR+) method is an intuitive, simple and conservative method that is introduced in ^16,34,35^ and is based on the concept that a significant phenotype can be called when the majority of the mutant mice lie outside the natural variation seen in the controls/WT. More extensive implementation of the RR+ is performed in the OpenStats that includes an open structure to make the comparison amongst all specified covariates as well as sub-levels. To overcome the misleading results from the data with repeated values, OpenStats reports empirical quantiles and adjust the threshold to the first distinct quantile.

## Comparison between analysis from OpenStats vs PhenStat

The data analysis pipeline in the IMPC requires several steps that are shown briefly in Figure 3. This consists of importing data to the operational environment and QC’ing before applying the statistical methods. The working datasets are formed carefully by splitting data based on predefined metadata to ensure all relevant information are packaged into a single dataset prior to applying the statistical analysis. The analysis engine is controlled by the statistical analysis software namely PhenStat or OpenStats. Ultimately, the statistical results are quality controlled for failures, errors and the agreement with the other available/manual implementations and if approved, passed to the downstream processes.

**Figure 3.**
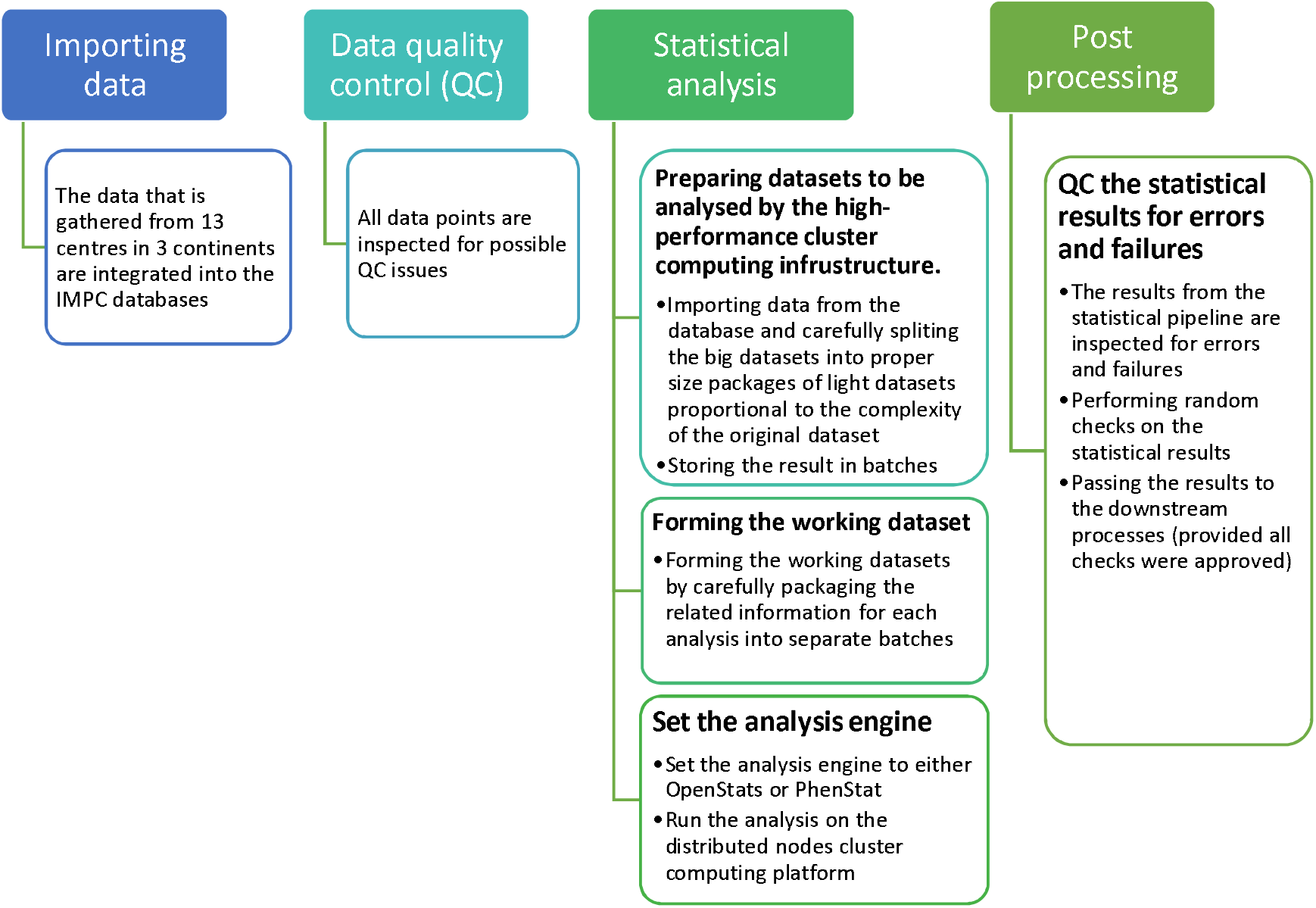
Schematic view of the IMPC statistical pipeline. The measurement of several parameters per specimen are collected from 13 centres all over the world, inspected for possible QC issues, carefully filtered to form individual working datasets, pre-optimised for being processed by the cluster computing platform and ultimately passed to the statistical analysis engine either PhenStat or OpenStats for the statistical analysis. The analysis engine is in charge of applying a proper statistical method to each working dataset and stores the analysis results in a format that enhances the downstream processes. All outputs from the statistical engine are inspected for the failures, errors and must pass a random QC check prior to being released to the downstream processes.

The comparison between the statistical analyses from OpenStats versus PhenStat is shown in Table 1 where two instances of the IMPC statistical pipeline equipped with OpenStats and PhenStat simultaneously ran on a cluster computing machine. The results consist of processing 226 distinct procedures (Supplementary Table S1) including 41 IMPC specific procedures such as IMPC calorimetry (IMPC_CAL), clinical blood chemistry (IMPC_CBC), haematology (IMPC_HEM), acoustic startle, pre-pulse inhibition (IMPC_ACS), insulin blood level (IMPC_INS) and body composition (IMPC_DXA), and over 3.8K parameters (1K+ IMPC parameters) (Supplementary Table S2). The complete list of IMPC procedures and parameters are available from www.mousephenotype.org/impress/pipelines. Because the analyses are performed on a farm of machines controlled by the cluster computing software, the direct measurement of time elapsed for each procedure/parameter is not straightforward. To alleviate this issue, the statistical pipeline reports the time spend per analysis/machine. Then, we report the average time for analysing the datasets in each procedure/parameter, which is normally the average of several hundred analyses. This ensures an unbiased estimation of the performance of the software.

**Table 1.**
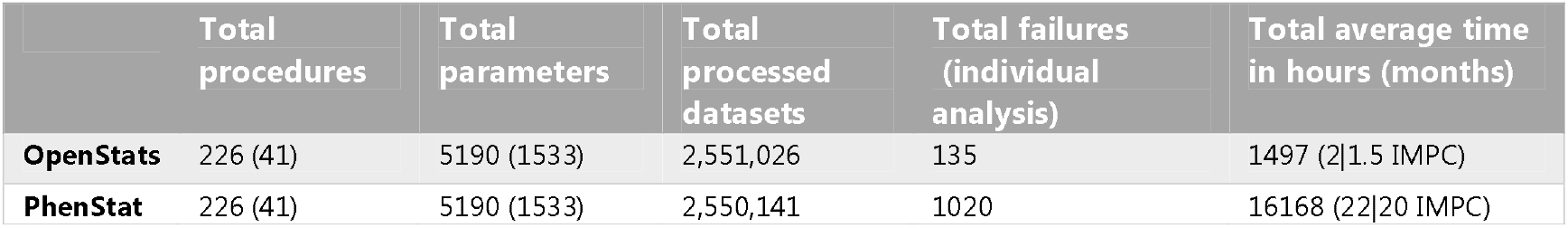
The comparison between OpenStats and PhenStat for analysing the IMPC continuous and categorical data. Each procedure contains several parameters that need to be analysed separately. Time elapse is estimated by adding up all spent times by carefully generated multi-processing jobs that equally fed into the OpenStats and PhenStat simultaneously. In all cases, the same procedure, parameter and data and analysis method are processed by OpenStats and PhenStat.

The analysis of the entire IMPC data using OpenStats and PhenStat on a hypothetical single-core machine takes a total of 24 months (~21.5 months for the IMPC specific procedures), including the categorical and continuous data. From that 24 months, 22 (92%) months (20 months for IMPC procedures) are accounted for by PhenStat and 2 (8%) months (1.5 months IMPC procedures) by OpenStats. Figure 4 (top row) shows the distribution of average time saved (in minutes) by utilising OpenStats software across the specific IMPC procedures (top ten) (left) and parameters (top 30 to save space) (right) whereas the bottom plots show the cases where PhenStat performs more efficient in time than OpenStats. These plots show a significant reduction in computational time from the OpenStats versus PhenStat software for processing the entire IMPC data. The full table of comparisons over all procedures is available in supplementary materials Table S3.

**Figure 4.**
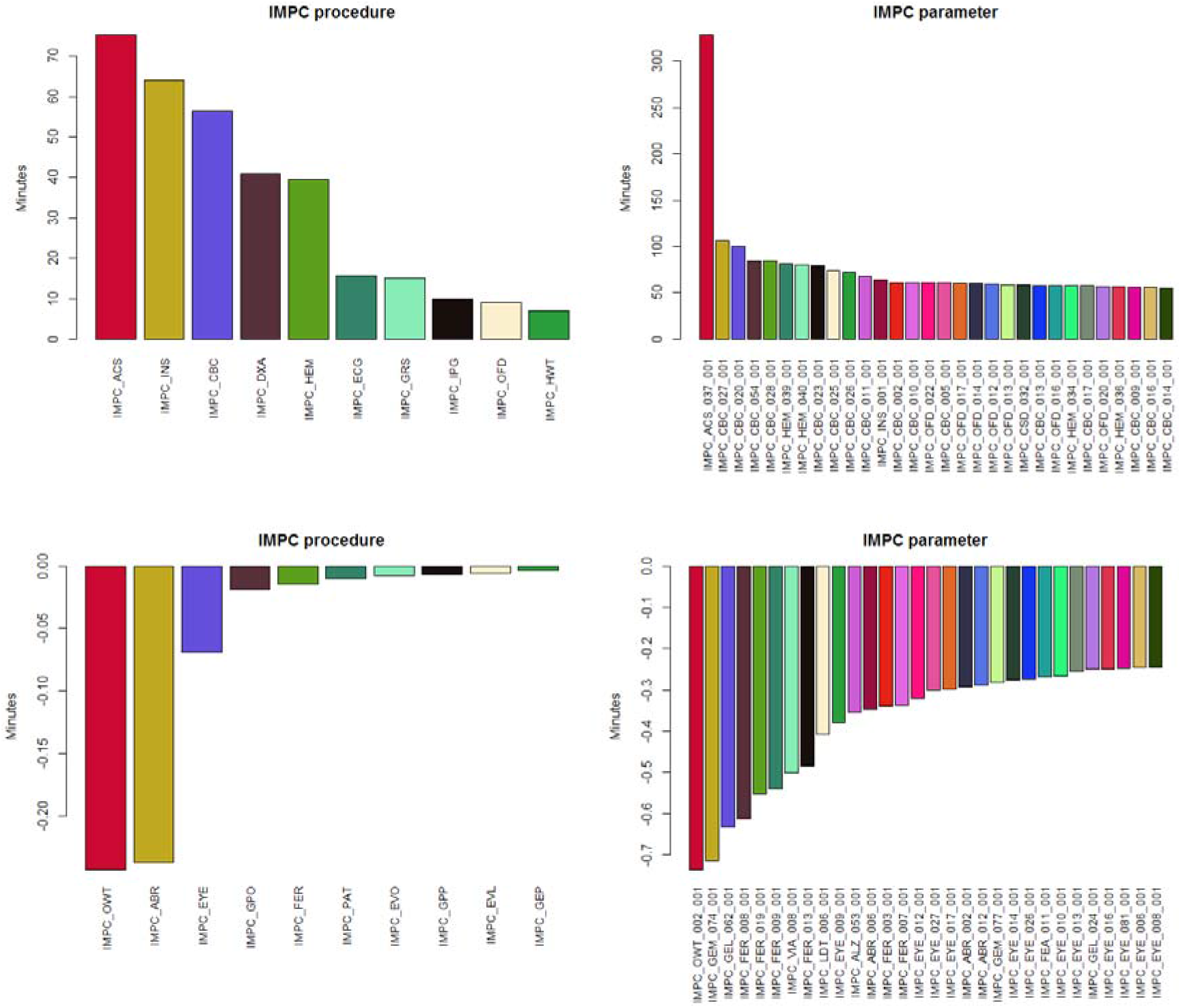
The comparison of the IMPC statistical pipeline analysed by OpenStats and PhenStat with respect to the time efficiency. (Top row) The left and right charts show the top (average) saving time in minutes by using OpenStats versus PhenStat over the IMPC procedures and parameters. The bottom row shows the top best (average) loses in minutes where PhenStat performs faster than OpenStats. These plots show that OpenStats improves the efficiency of the IMPC statistical pipeline.

We further performed a confirmatory analysis to compare the agreement between the statistical results, Genotype effect p-value in particular, from PhenStat and OpenStats. Our results show an overall agreement of 99% between the two software, 99.9%, 99.9%, and 98.9% for Fisher’s exact test, Reference Range plus, and Linear Mixed model frameworks respectively. The main cause of the disagreement between the two software is using Monte Carlo simulation-based method for the Fisher’s exact test and Reference Range plus as well as new model selection strategy based on Akaike information criterion for small samples (AICc) for the internal optimisation of the software and the natural diversity in the data that sometimes violates the assumptions of the applied model for the Linear Mixed model frameworks. For instance, https://bit.ly/2WxI2Io is an example dataset from IMPC Acoustic Startle and Pre-pulse Inhibition (PPI) procedure. This data is right-skewed and should be considered carefully. In this example, the internal optimisation of PhenStat removes the effect of the bodyweight (at the level of 0.05) as a covariate in the linear mixed model under the settings in Eq. (1) whereas OpenStats preserves it in the model. The consequence is a 2-fold decrease in the magnitude of the genotype effect p-value, 0.49 in PhenStat and 0.18 in OpenStats. We should stress that there is no right answer in this example as there is a violation of the model assumptions/residuals. The next example in https://bit.ly/2YB4int represents a dataset from the IMPC Haematology procedure and depicts the inconsistency in the statistical results that is a product of the missing values in body weight (14% in total) and the outliers in response (5% based on Tukey’s criteria with k = 3). In this case, PhenStat omits the bodyweight effect from the linear mixed model in Eq. (1) (Genotype effect p-value 0.0004) whereas OpenStats keeps the body weight in the model but excludes the missing values from the analysis (OpenStats Genotype effect p-value 0.97). We should stress that regardless of the statistical software, unusual cases should be interpreted carefully.

## Discussion

Establishing precise, robust, reliable, and reproducible statistical pipelines for high-throughput phenotyping screens is challenging and is the subject of recent research topics ^16,19,27,28,36–38^. With ever-growing phenotypic data, more consideration is required for scalability and versatility of the statistical pipeline to model different possible scenarios as efficiently as possible. One challenge the International Mouse Phenotyping Consortium (IMPC) faces is the diversity in data that demands more complex statistical methods with minimal latency. Here we introduced OpenStats, a freely available R package for systematically analysing high-throughput phenotypic data. The R package allows a fully customised analysis plan for the implemented methods namely: linear mixed model, Fisher’s exact test and Reference Range plus, as well as a comprehensive workflow with a focus on simplicity, efficiency, scalability and completeness that offers more than the raw statistical results and more than the counterparts in the literature. The performance of the new software compared to the current implementation of the statistical pipeline in IMPC is assessed on more than 61M data points and 4M + analyses. Our comparisons show on average 90% reduction in time spent by adopting the new software while 99% of the results remain similar between OpenStats and the closest counterpart, PhenStat, for the IMPC data. The speed efficiency of OpenStats lies in the fact that the software utilises a “start-update” strategy instead of “start-terminate”. That is, each model is formed by updating the previous one in contrast to fitting a brand-new model at each step. Besides the advantages of the software, there are a number of limitations to OpenStats including dependency to the statistical pipelines built on R and difficulties with exchanging the statistical methods with other statistical analysis software such as Python.

OpenStats addresses other challenges beyond the need to scale analysis to large datasets. One example is irreproducibility in results from animal experiments that is cited as a major contributor to explain many drugs failure in the development pipeline ^39^. To address this concern, the ARRIVE (Animal Research: Reporting of *In Vivo* Experiments) guidelines have recently been updated to emphasis reporting on statistical methods including clearly stating the statistical method that is used and whether the data meet the assumptions of the statistical approach ^40^. As part of its operation, OpenStats assesses whether data fits the requirements for a statistical test, such as not having enough data for performing interaction tests in the linear mixed model or assessing the normality of input data/model residuals. OpenStats also provides the visualisation tools required to diagnose the fitted model. The resulting analyses are clearly defined by which method was used, promoting reproducibility and repeatability of the results and the statistical models.

Increasingly researchers and stakeholders such as funders are demanding that biological research data and their analyses follow the FAIR principles - Findable, Accessible, Interoperable, and Reusable ^41^. OpenStats contributes to FAIR data by assessing input data for completeness, redundancy, and any mismatch in variable format and/or labels. It also provides semantic normalisation such as sex, Sex and gender to a unified term “Sex” for common biological data variables in order to promote interoperability and reusability of data. Critically, OpenStats enables reusability of statistical methods by being freely available from the well-known BioConductor software project (www.bioconductor.org/packages/OpenStats) allowing any researcher to reproduce and reuse analyses from others’ research while ensuring their own analysis is FAIR.

In summary, OpenStats promotes FAIR data and better reproducibility of biological research results while providing a means to scale to the larger and more complex datasets being generated by the research community.

## Future study

Future work could be deriving a quality score that represents the quality of individual statistical analysis from the high-throughput genomic pipelines by testing the different aspects of the input data and the analysis results for the specified analysis framework.

## Funding

This work was supported by the NIH Common Fund UM1-HG006370.

## Notes

https://bioconductor.org/packages/release/bioc/html/OpenStats.html

### Competing Interest Statement

The authors have declared no competing interest.

## References

1. Prinz, F., Schlange, T. & Asadullah, K. Believe it or not: how much can we rely on published data on potential drug targets? Nat. Rev. Drug Discov. 10, 712–712 (2011).

2. Collins, F. S. & Tabak, L. A. NIH plans to enhance reproducibility. Nature 505, 612–613 (2014).

3. Kilkenny, C., Browne, W. J., Cuthill, I. C., Emerson, M. & Altman, D. G. Improving bioscience research reporting: The arrive guidelines for reporting animal research. Animals 4, 35–44 (2013).

4. Goktug, A. N., Ong, S. S. & Chen, T. GUItars: A GUI Tool for Analysis of High-Throughput RNA Interference Screening Data. PLoS One 7, (2012).

5. Schulz, J. B., Cookson, M. R. & Hausmann, L. The impact of fraudulent and irreproducible data to the translational research crisis – solutions and implementation. J. Neurochem. 139, 253–270 (2016).

6. Holmes, S. Statistical proof? The problem of irreproducibility. Bull. Am. Math. Soc. 55, 31–55 (2018).

7. Karp, N. A. et al. Applying the ARRIVE Guidelines to an In Vivo Database. 13, e1002151 (2015).

8. Ozonoff, D. M. & Grandjean, P. What is useful research? The good, the bad, and the stable. Environ. Heal. A Glob. Access Sci. Source 19, (2020).

9. Hirsch, C. & Schildknecht, S. In vitro research reproducibility: Keeping up high standards. Frontiers in Pharmacology vol. 10 (2019).

10. Koscielny, G. et al. The International Mouse Phenotyping Consortium Web Portal, a unified point of access for knockout mice and related phenotyping data. Nucleic Acids Res. 42, D802–D809 (2014).

11. Brown, S. D. M. & Moore, M. W. The International Mouse Phenotyping Consortium: Past and future perspectives on mouse phenotyping. Mamm. Genome 23, 632–640 (2012).

12. Bradley, A. et al. The mammalian gene function resource: The International Knockout Mouse Consortium. Mamm. Genome 23, 580–586 (2012).

13. De Angelis, M. H. et al. Analysis of mammalian gene function through broad-based phenotypic screens across a consortium of mouse clinics. Nat. Genet. 47, 969–978 (2015).

14. Kurbatova, N., Karp, N., Mason, J. & Haselimashhadi, H. PhenStat : statistical analysis of phenotypic data. Bioc.Ism.Ac.Jp 1–9 (2016).

15. R Team Core. . R Foundation for Statistical Computing, Vienna, Austria. 2019 (2019).

16. Kurbatova, N., Mason, J. C., Morgan, H., Meehan, T. F. & Karp, N. A. PhenStat a tool kit for standardized analysis of high throughput phenotypic data. PLoS One 10, e0131274 (2015).

17. Gilbert, G. E. Linear Mixed Models: A Practical Guide Using Statistical Software. J. Am. Stat. Assoc. 103, 427–428 (2009).

18. De Angelis, M. H. et al. Analysis of mammalian gene function through broad-based phenotypic screens across a consortium of mouse clinics. Nat. Genet. 47, 969–978 (2015).

19. Haselimashhadi, H. et al. Soft Windowing Application to Improve Analysis of High-throughput Phenotyping Data. Bioinformatics (2019) doi:10.1093/bioinformatics/btz744.

20. Kurbatova, N., Mason, J. C., Morgan, H., Meehan, T. F. & Karp, N. A. PhenStat: A Tool Kit for Standardized Analysis of High Throughput Phenotypic Data. PLoS One 10, e0131274 (2015).

21. Dennis, B., Ponciano, J. M., Taper, M. L. & Lele, S. R. Errors in Statistical Inference Under Model Misspecification: Evidence, Hypothesis Testing, and AIC. Front. Ecol. Evol. 7, 372 (2019).

22. Patefield, W. M. Algorithm AS 159: An Efficient Method of Generating Random R × C Tables with Given Row and Column Totals. Appl. Stat. 30, 91 (2006).

23. Fisher, R. A. The Logic of Inductive Inference. J. R. Stat. Soc. 98, 39 (2006).

24. Clarkson, D. B., Fan, Y. & Joe, H. A remark on algorithm 643: FEXACT: an algorithm for performing Fisher’s exact test in r x c contingency tables. ACM Trans. Math. Softw. 19, 484–488 (1993).

25. Agresti, A. Categorical data analysis. (Wiley, 2003).

26. Pinheiro, J. C. & Bates, D. M. Mixed-effects models in S and S-PLUS. (Springer, 2000).

27. Karp, N. A. et al. Impact of temporal variation on design and analysis of mouse knockout phenotyping studies. PLoS One 9, e111239 (2014).

28. Karp, N. A., Melvin, D. & Mott, R. F. Robust and Sensitive Analysis of Mouse Knockout Phenotypes. PLoS One 7, e52410 (2012).

29. Suárez, E., Pérez, C. M., Rivera, R. & Martínez, M. N. Applications of Regression Models in Epidemiology. Applications of Regression Models in Epidemiology (John Wiley & Sons, Inc., 2017). doi:10.1002/9781119212515.

30. Burnham, K. P., Anderson, D. R. & Losos, J. B. Model selection and multimodel inference. A practical information-theoretical approach. Ecology Letters vol. 11 (Springer, 2002).

31. Cohen, J. Statistical Power Analysis for the Behavioral Sciences. Statistical Power Analysis for the Behavioral Sciences https://books.google.co.uk/books?id=rEe0BQAAQBAJ&printsec=frontcover&dq=coheneffectsize&hl=en&sa=X&ved=0ahUKEwj51ezztpfnAhUaHMAKHXQNCtQQ6AEIOjAC#v=onepage&q=coheneffectsize&f=false (2013) doi:10.4324/9780203771587.

32. Royston, J. P. An Extension of Shapiro and Wilk’s W Test for Normality to Large Samples. Appl. Stat. 31, 115 (2006).

33. Conover, W. J. & Conover, W. J. Practical Nonparametric Statistics (Wiley Series in Probability and Statistics). (John Wiley & Sons, 1980).

34. White, J. K. et al. XGenome-wide generation and systematic phenotyping of knockout mice reveals new roles for many genes. Cell 154, 452 (2013).

35. Cook, M. N. et al. Neurobehavioral mutants identified in an ENU-mutagenesis project. Mamm. Genome 18, 559–572 (2007).

36. Willis, R. Must try harder. Community Care 483, 32–33 (2006).

37. Begley, C. G. & Ellis, L. M. Drug development: Raise standards for preclinical cancer research. Nature 483, 531–533 (2012).

38. Baker, D., Lidster, K., Sottomayor, A. & Amor, S. Two Years Later: Journals Are Not Yet Enforcing the ARRIVE Guidelines on Reporting Standards for Pre-Clinical Animal Studies. PLoS Biol. 12, e1001756 (2014).

39. Freedman, L. P., Cockburn, I. M. & Simcoe, T. S. The Economics of Reproducibility in Preclinical Research. PLOS Biol. 13, e1002165 (2015).

40. Sert, N. P. du et al. The ARRIVE guidelines 2019: updated guidelines for reporting animal research. bioRxiv 03181 (2019) doi:10.1101/703181.

41. Wilkinson, M. D. et al. Evaluating FAIR maturity through a scalable, automated, community-governed framework. Sci. data 6, 174 (2019).

